# Human Intestinal Absorption Revisited: The 1998 Descriptors Survive, and the Method Extends to Permeability

**DOI:** 10.64898/2026.10.03.756481

**Authors:** Steven M. Muskal, Matthew D. Wessel

**Affiliations:** Eidogen-Sertanty, Inc., Oceanside, CA 92056; Thermo Fisher Scientific, Sisters, OR 97759

**Keywords:** human intestinal absorption, QSPR, genetic algorithm, descriptor selection, cyclic peptides, membrane permeability, oral bioavailability, repeated extraction

## Abstract

In 1998 we predicted percent human intestinal absorption (HIA) for 86 drug and drug-like compounds from six molecular descriptors chosen by a genetic algorithm with a small neural network serving as the fitness function. That result rested on one division of the compounds into three sets: training, cross-validation and external prediction. Repeating the division shows it was a rather fortunate one. An extraction is one random draw from a wide distribution, so we repeat every experiment 250 times and report the result as a distribution. Across 250 stratified extractions, 14 match the published cross-validation error and only one matches the published external prediction error. Reconstructed this way, the published six descriptors give a mean extraction error roughly twice the external prediction value we reported in 1998 (Table 1). We then applied the same six descriptors to a larger set of 780 compounds from a public compilation. Under the 1998 network they give 22.50 %HIA units on training compounds against 25.77 on extracted compounds, whereas on the 86 they give 13.43 against 30.39. More compounds close the gap between fitting and prediction. Four datasets are modeled here: 780 small molecules with HIA measured in man, 3,771 small molecules and 1,281 cyclic peptides measured across Caco-2 cell monolayers, and 7,298 cyclic peptides measured across artificial membranes. Each is its own prediction problem, and one procedure is applied to all four. A genetic algorithm chooses which descriptors to use and how many, and a random forest grades each choice on compounds its own trees never saw. The descriptors chosen most often across the 250 extractions form one consensus set for each dataset. For absorption, 25 consensus descriptors under a random forest reach an extraction rms of 20.47 %HIA units at R 0.76. On the same compounds and the same extractions, the six 1998 descriptors under the 1998 network reach 25.77 at R 0.59. The four models are served at predicthia.ai, where a structure can be drawn or pasted, scored against its training set for similarity, and the training data with fitted models downloaded. Each model served is its fixed consensus set fitted on every compound in its dataset; the accuracy quoted beside a prediction comes from evaluating that same set across 250 extractions (Table 3). The fixed sets hold 25 descriptors for absorption, 55 for Caco-2 small-molecule permeability, 50 for Caco-2 peptide permeability and 25 for artificial-membrane permeability.

**Graphical abstract:** The search, in the layout of the original study. Left, the genetic algorithm cycle, with fitness now the out-of-bag error of a random forest. Middle, four candidate descriptor sets drawn at their own lengths, one cell per descriptor, each scored by its own forest, and below them a crossover worked through as in 1998. Right, the distribution of extraction error across 250 extractions of the 780 absorption compounds, the right panel of Figure 1, and the deployed 25-descriptor model against measurement.

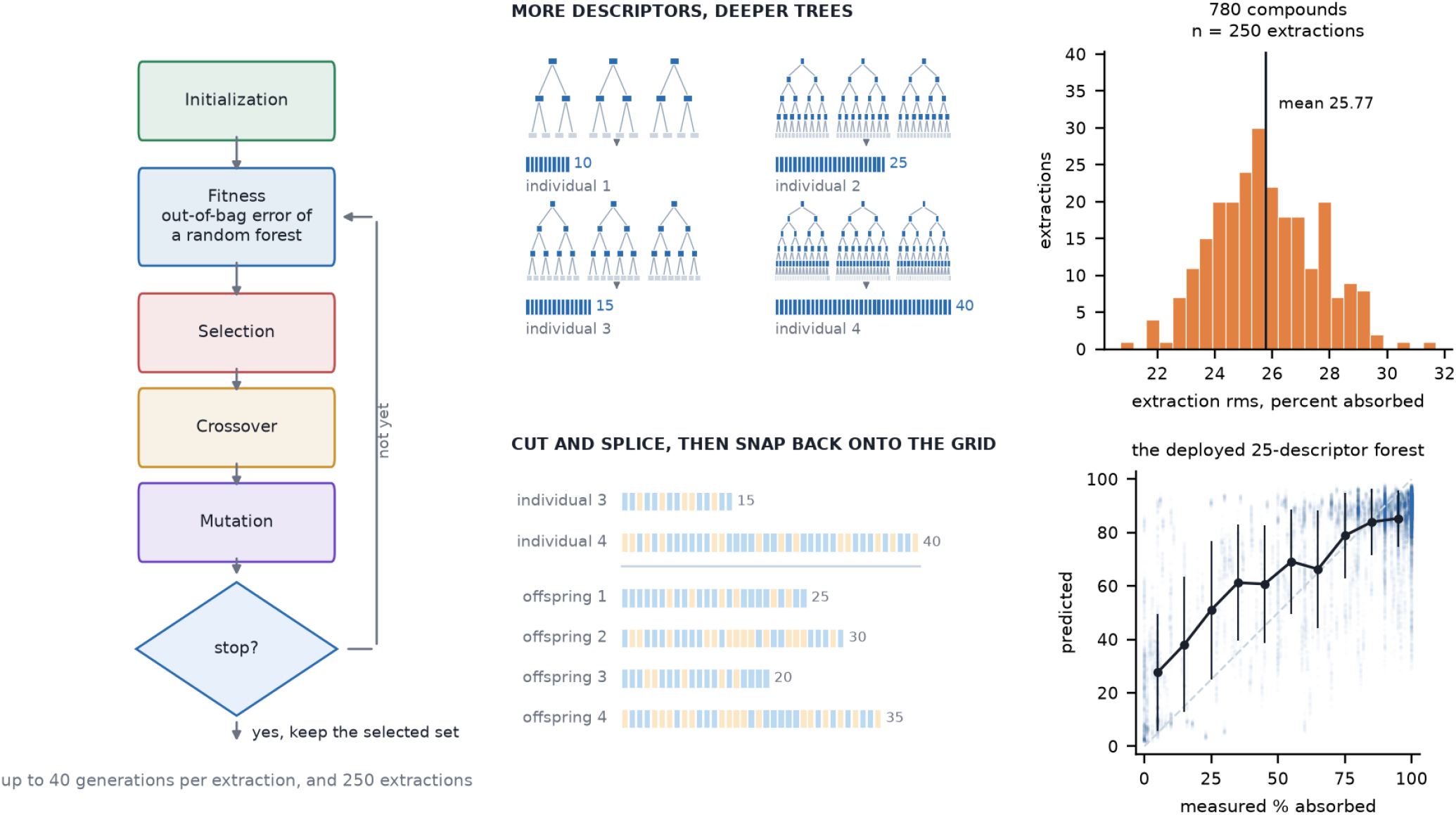

## 1. Introduction

Human intestinal absorption is a property of a compound that can be estimated while its structure is still being designed. It measures passage from an oral dose across the intestinal wall, typically in the duodenum. Subsequent first-pass metabolism determines how much of the absorbed dose reaches systemic circulation, so percent absorbed and oral bioavailability are distinct endpoints and should not be confused.

In 1998 we modeled percent human intestinal absorption (HIA) for 86 drug and drug-like compounds using six molecular descriptors selected by a genetic algorithm and a small neural network.[1] The paper reported rms errors of 9.4 %HIA units for training, 19.7 for cross-validation and 16.0 for ten external prediction compounds. That paper has since been cited more than 500 times as of October 2026, and the endpoint has been modeled repeatedly on progressively larger and more diverse compound sets. The physical basis of such models developed alongside our original work: polar surface area,[2] Lipinski’s rule of five,[3] and molecular flexibility.[4] Later compilations expanded the absorption data,[5,6] while analyses of training-set composition and imbalance showed why an accuracy estimate depends on which compounds are evaluated.[7,8] Repeated modeling on a common dataset provided a way to examine that dependence.[9]

We return first to the six descriptors published in 1998. We evaluate the reconstructed network over 250 stratified extractions of its original 86 compounds, then carry those descriptors unchanged to a larger absorption set of 780 compounds. This separates the behavior of the published descriptor set from the performance of one favorable draw.

We then apply one descriptor-selection and deployment procedure to four endpoints: percent human intestinal absorption, small-molecule and peptide permeability across Caco-2 monolayers, and peptide permeability across artificial membranes. CycPeptMPDB supplies the peptide measurements.[10] The two permeability assays are kept separate because each defines its own prediction target. For each endpoint, a genetic algorithm search is repeated over 250 extractions, the descriptors chosen most often form one fixed consensus set, and that set is evaluated over the same extractions before a model is fitted on every compound for predicthia.ai.

## 2. Methods

### 2.1 Data

Absorption was taken from a public compilation of human intestinal absorption measurements.[11] Structures were taken from the PubChem identifier the compilation supplies for each row. Two entries for a fluorescent paracellular permeability marker were excluded, and one compound whose geometry would not embed was dropped, leaving 780 compounds.

Where several absorption measurements were available for a compound, their median was used. The original 86 compounds and their absorption values were taken from Table 1 of the 1998 study.[1]

**Table 1.** The 1998 procedure, as published and as repeated. All three blocks use the same six descriptors and the same model, the 6-4-1 logistic feed-forward network of the 1998 paper. The first block is that paper’s own result on its 86 compounds, from the single division of those compounds it reported. The second block is a reconstruction of that procedure on the same 86 compounds, repeated over 250 stratified extractions. The group sizes are 62, 11 and 13 against the original 67, 9 and 10, and the descriptors were reimplemented from their published definitions. The third carries the same six descriptors, unchanged, onto 780 absorption compounds, also over 250 extractions. Errors are in %HIA units: rms is root mean square error and mae is mean absolute error, each a mean over the extractions with its standard deviation in parentheses. R is the Pearson correlation on the extracted group. The published block has no standard deviations because it is one split, and its R was computed over all 86 compounds including the 67 used for training. The 1998 paper also reported a correlation of 0.93, computed over all 86 compounds including the 67 used for training. It is not an external prediction statistic and is not shown in the table.

| Dataset | Group | rms | mae | R |
| --- | --- | --- | --- | --- |
| 1998, as published, one split | Training | 9.40 | 6.70 |  |
|  | Cross-validation | 19.70 | 15.40 |  |
|  | External prediction, ten compounds | 16.00 | 11.00 |  |
| 86 compounds, 250 extractions | Training | 13.43 (1.90) | 9.46 (1.30) |  |
|  | Cross-validation | 29.37 (5.98) | 22.03 (4.65) |  |
|  | Extracted | 30.39 (5.16) | 22.63 (4.26) | 0.46 (0.23) |
| 780 compounds, 250 extractions | Training | 22.50 (0.50) | 16.38 (0.42) |  |
|  | Cross-validation | 25.56 (2.05) | 18.68 (1.45) |  |
|  | Extracted | 25.77 (1.82) | 18.76 (1.16) | 0.59 (0.07) |

**Table 2.** The six descriptors published in 1998 and the similar metric the absorption search selects, over 250 extractions of the 780 compound

| The 1998 descriptor, and what it measures | Similar metric identified by the modern search | Chosen in |
| --- | --- | --- |
| CHDH-1, charge carried on donatable hydrogens | mean partial charge over the molecule | 100.00% |
| SAAA-2, acceptor surface area per acceptor | topological polar surface area | 35.60% |
| SCAA-2, acceptor surface weighted by charge | partial-charge weighted surface area | 32.00% |
| NSB, number of single bonds | rotatable bond count | 26.40% |
| GRAV-3, cube root of the gravitational index, the spread of atomic mass in three dimensions | radius of gyration | 11.60% |
| SHDW-6, normalized projection of the molecule on a plane | sphericity index | 4.00% |

**Table 3.** The four deployed models. Every value is measured on the extracted group, as the mean over 250 extractions of the fixed consensus set, with standard deviations in parentheses. Errors are in %HIA units for absorption and log10 Papp units for the three permeability models. The Caco-2 peptides were searched on a grid extending to 60 descriptors and the Caco-2 small molecules on one extending to 55.

| Endpoint | Compounds | Descriptors | rms | mae | R |
| --- | --- | --- | --- | --- | --- |
| Absorption, % absorbed | 780 | 25 | 20.47 (1.71) | 14.79 (1.11) | 0.76 (0.05) |
| Caco-2 small molecules, log<sub>10</sub> Papp | 3,771 | 55 | 0.35 (0.01) | 0.26 (0.01) | 0.84 (0.01) |
| Caco-2 peptides, log<sub>10</sub> Papp | 1,281 | 50 | 0.49 (0.04) | 0.34 (0.02) | 0.84 (0.03) |
| Artificial-membrane peptides, log<sub>10</sub> Papp | 7,298 | 25 | 0.82 (0.03) | 0.49 (0.02) | 0.67 (0.02) |

Peptide permeability was taken from CycPeptMPDB[10] and divided into two sets that are not interchangeable: 1,281 peptides with Caco-2 monolayer measurements and 7,298 with artificial-membrane measurements. Permeability is reported as log10 Papp, with Papp in cm/s. Both sets record permeability below the detection limit as exactly -10.00, for 21 peptides in the Caco-2 set and 269 in the artificial-membrane set. These values are placeholders and were not measured. These compounds are included in training and in the reported errors. They are omitted only from Figure 5, where plotting them against a measured axis would be misleading. The two sets share 588 of the 1,281 Caco-2 peptides. Where a single study measured a peptide in both assays, on 229 peptides, the two assays correlate 0.62 with the artificial-membrane value about half a log unit below the Caco-2 value. Comparisons drawn across different studies carry no relationship and are not used here. The absorption set shares one compound with the peptide sets.

A fourth dataset extends the procedure to small-molecule permeability in the same assay as the Caco-2 peptides. The OpenADMET consortium released the full post-challenge dataset of the OpenADMET-ExpansionRx blind challenge, lead optimization measurements contributed by Expansion Therapeutics across 7,608 compounds and nine endpoints.[25,26] Of those, 3,773 carry an apparent permeability apical to basolateral across a Caco-2 monolayer, reported as Papp in 10-6 cm/s. Two values recorded as exactly zero were dropped, since the logarithm is undefined, leaving 3,771 compounds. Permeability was converted to log10 Papp with Papp in cm/s, the same quantity and units as the peptide models, giving a range of -8.00 to -4.29. The release also carries a Caco-2 efflux ratio, which reports active transport rather than permeability and is not modeled here. The released files carry only in-range measurements; the accompanying raw file holds 33 further values marked with a less-than modifier, each at its own threshold rather than at one detection limit, and those are not used. This differs from the peptide sets, where values below the detection limit share a single placeholder at -10.00.

### 2.2 Descriptors

Descriptors were computed under RDKit 2025.09.5, and the deployed models are pinned to that release because descriptor values depend on it.[12] The absorption pool contains 1,301 descriptors from charge, charged partial surface area,[13] two-dimensional and three-dimensional RDKit blocks, Morgan bits,[14] the six descriptors published in 1998, and a block of size and composition terms: molecular weight, heavy atom and bond counts, rotatable bonds, ring count and element counts. The peptide pool contains 1,257 descriptors from Morgan bits, two-dimensional RDKit descriptors and a peptide-specific block. The Caco-2 small molecules use the same 1,301-descriptor pool as absorption. Any descriptor holding a non-finite value on any compound was dropped whole; nothing was imputed and no value was filled.

The six published descriptors were reimplemented from their definitions: single-bond count, normalized shadow projection,[15] charge on donatable hydrogens, two hydrogen-bond acceptor surface terms and the cube root of the gravitational index. For the shadow descriptor, atoms were projected as van der Waals discs after alignment to the principal axes of inertia. Geometries were obtained by distance-geometry embedding[16,17] and force-field optimization,[18] and empirical partial charges were used.[19] These six descriptors were fixed in both reproduction experiments.

### 2.3 Extraction

An extraction is one stratified random draw of the compounds into a training group and an extracted group, stratified by absorption band or by decile of permeability. Fifteen percent of the compounds form the extracted group. The random forest experiments need no separate cross-validation group, since out-of-bag scoring requires none, and those compounds remain in training. Every experiment reported here uses 250 extractions. The seed of each extraction is 1000 plus its index.

Each extraction draws its groups at random, so extractions will overlap: a compound in the extracted group of one extraction is likely to be in the training group of another. Over 250 extractions, each compound falls in the extracted group about 38 times, so the reported error is an average over many divisions of the compounds. The count of 250 was chosen from the spread of the single-extraction result. On the 86 compounds, extraction error ranges from 13.73 to 45.81 across extractions, and 250 extractions bring the standard error of the mean below a third of a %HIA unit, which is small against the differences reported here.

Objective selection, a screen that reads only the descriptors and never the measurement, was applied inside each extraction on its training compounds only: any descriptor with more than 80.00 percent identical values was dropped, then one of any pair correlating above 0.90. Between 127 and 184 descriptors remain, depending on the endpoint, with medians over the 250 extractions of 148 for absorption, 181 for the Caco-2 small molecules, 142 for the Caco-2 peptides and 131 for the artificial-membrane peptides. In the reproduction experiments, the published six descriptors were retained without a new selection.

The network reproduction includes a separate cross-validation group. Its group sizes are 62 training, 11 cross-validation and 13 extracted on the original 86 compounds, and 584, 79 and 117 on the larger absorption set.

The random forest extractions need no cross-validation group. Their training and extracted sizes are:

- absorption, 663 training and 117 extracted
- Caco-2 small molecules, 3,203 training and 568 extracted
- Caco-2 peptides, 1,089 and 192
- artificial-membrane peptides, 6,204 and 1,094

### 2.4 Descriptor selection

Descriptor sets were selected by a genetic algorithm.[20,21] Each candidate is a set of descriptors, and the number of descriptors in a set is itself searched, on a grid of 5, 10, 15, 20, 25, 30, 35 and 40 within a single run. The Caco-2 peptide search extends this grid to 60 descriptors in the same steps of five. The count is neither fixed in advance nor tuned by hand. A population of 60 candidate sets was evolved for up to 40 generations. In crossover, each parent was cut at its own point, the head of one was joined to the tail of the other, and the size of the child was rounded to the nearest count on the grid. Where crossover left a child short of that count, the shortfall was drawn from the parents, and a handful of sets finished one or two descriptors below a grid value. Those extractions are counted at the size they reached, which is why Figure 2 carries a small number of bars off the grid. In mutation, a set was enlarged by one grid step, reduced by one grid step, or changed by exchanging one descriptor for another. The Caco-2 small molecules were searched on a grid of 5 to 55 descriptors in steps of five, using the small-molecule descriptor pool rather than the peptide pool.

**Figure 1.**
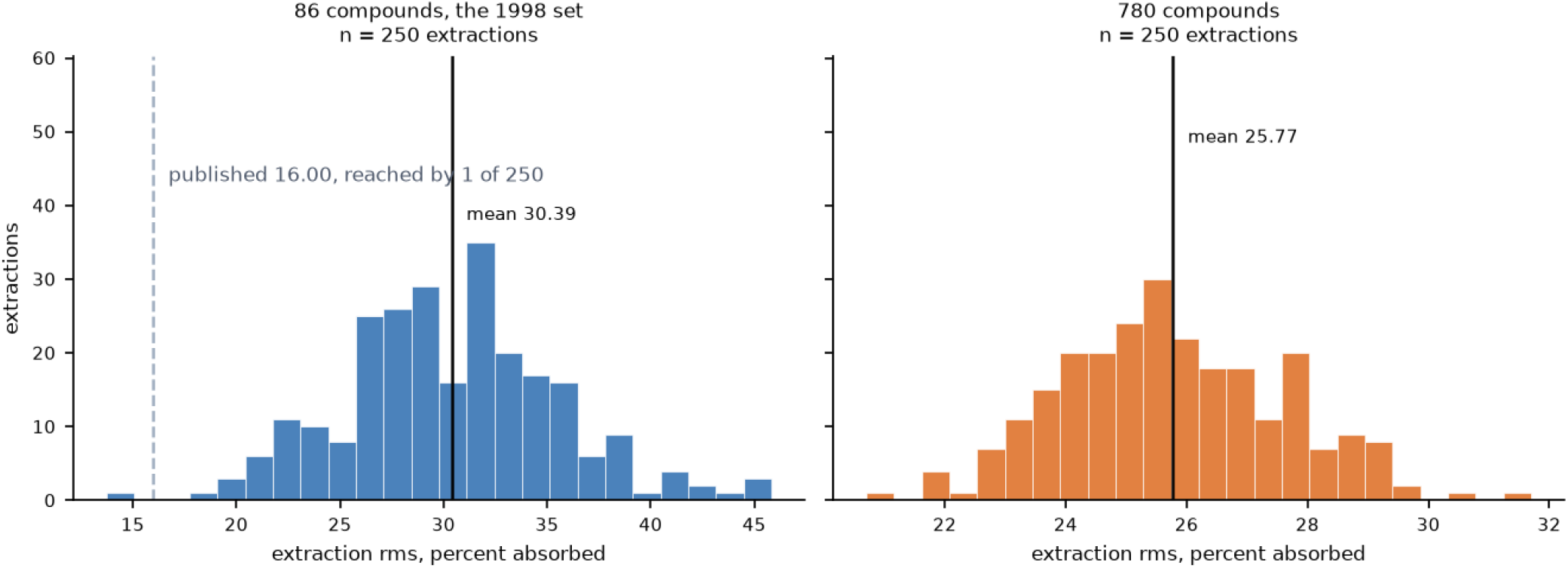
Distribution of extraction rms over 250 stratified extractions using the published six descriptors and the 1998 network. Left, the original 86 compounds. Right, the larger absorption set of 780 compounds. The dashed line marks the published external prediction rms of 16.00 %HIA units; solid lines mark the means.

**Figure 2.**
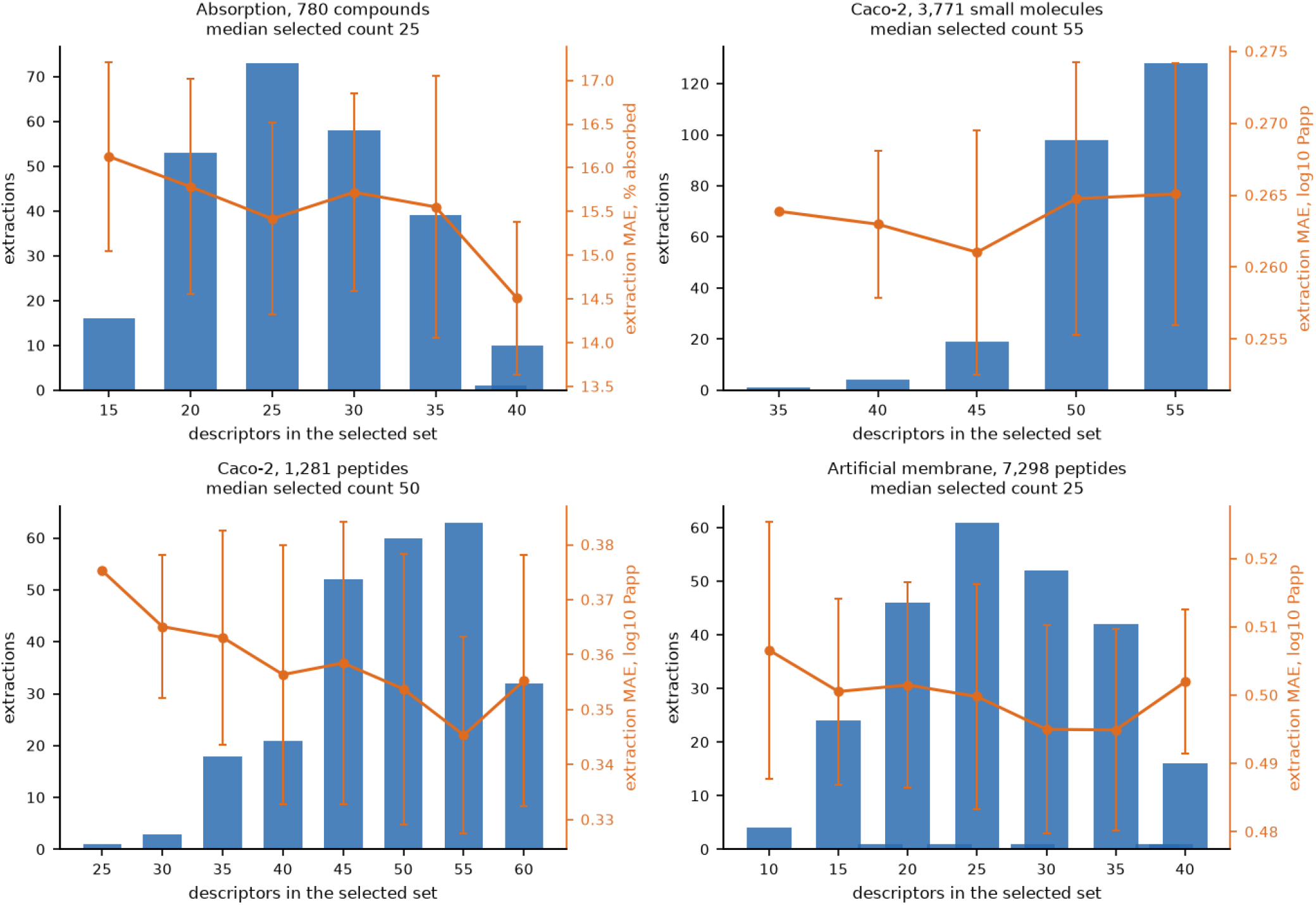
How many descriptors the search chooses, and what that costs. Bars, left axis: the number of the 250 extractions whose selected descriptor set had that many descriptors. Orange line, right axis: the mean extraction error of those extractions at each count, with one standard deviation. Error is mae, the mean absolute error, which is the average size of the miss in the units of the measurement, % absorbed on the top left panel and log10 Papp on the other three. Top left, absorption. Top right, the Caco-2 small molecules. Bottom left, the Caco-2 peptides. Bottom right, the artificial-membrane peptides. The number of descriptors was searched on a grid of 5 to 40 in steps of five, of 5 to 55 for the Caco-2 small molecules, and of 5 to 60 for the Caco-2 peptides; the few bars off that grid are extractions where crossover left a set one or two descriptors short. Absorption and the artificial-membrane peptides settle at a median of 25 descriptors, the Caco-2 small molecules at 55 and the Caco-2 peptides at 50, while extraction error changes little across each range.

The fitness of a candidate set was the out-of-bag error of a random forest fitted on the training compounds using that set’s descriptors. Each tree was grown on a bootstrap sample, and predictions for a training compound were obtained only from trees whose bootstrap samples excluded it. Roughly a third of the training compounds are absent from any given tree. Nothing outside the training group was consulted while descriptors were being chosen, and the extracted group was read once per extraction after the search had ended. The genetic algorithm selects the descriptors; the forest only grades the selection. A forest fitted on the whole pool would require every descriptor in it for every new compound, which is why the selected set of 25 to 55 is what we compute and publish.

Choosing descriptors by minimizing error is itself a form of fitting, so the error observed during a search cannot serve as an independent estimate of accuracy.[22,23] The independent estimate comes from the extracted group of each extraction, which the search never reads.

### 2.5 Models

The 1998 reproduction used the architecture of the 1998 paper, a 6-4-1 logistic feed-forward network with six inputs, four hidden units and one output. The same architecture was used on the 86 compounds and on the 780. The deployed models were fitted with random forests of 300 trees and a minimum of two compounds per leaf.

The random forest was chosen for the deployed models for three practical reasons. It has no architecture to size, so no hidden layer has to be chosen for each candidate descriptor set, and a search over descriptor sets does not become a search over architectures at the same time. It fits quickly enough to score a population of sixty descriptor sets over forty generations inside each of 250 extractions. And it supplies an out-of-bag error, which gives the search an objective computed entirely inside the training group. The purpose of the reproduction experiments is to repeat the published procedure, so the 1998 network is kept there. Each evaluation of a fixed consensus set averaged eight random forests grown from different seeds, while the deployed model is a single forest at seed 0. Conformers were generated with the ETKDGv3 distance-geometry generator from a fixed seed, with ten conformers optimized using MMFF and the lowest-energy conformer retained. The seed is part of the method because changing it changes the three-dimensional descriptors. Both models were implemented with scikit-learn.[24]

### 2.6 From 250 extractions to a deployed model

One selected descriptor set was obtained from each of the 250 extractions. The consensus set was formed from the descriptors chosen most often, with its size set to the median size of the selected sets, 25 descriptors for absorption, 55 for the Caco-2 small molecules, 50 for the Caco-2 peptides and 25 for the artificial-membrane peptides. Because the consensus set is assembled across extractions and is not itself one of the selected sets, its error was measured directly. The consensus set was held fixed and evaluated over the same 250 extractions with no further selection, and those are the only statistics reported for a deployed model.

Each deployed model was then fitted on every compound in its dataset. A deployed model has no extracted group, and the extraction statistics state its expected performance. Errors are reported as rms and mae, with standard deviations across the 250 extractions, and R is the Pearson correlation on the extracted group. Absorption errors are in %HIA units and permeability errors in log10 Papp units.

## 3. Results

### 3.1 The published six descriptors over repeated extractions

The published six descriptors were applied without a new descriptor search to the original 86 compounds and to the larger absorption set. The model remained the 1998 network in both experiments. Every reported value in Table 1 is a mean over 250 stratified extractions.

On the 86 compounds, training rms is 13.43 %HIA units, cross-validation rms is 29.37 and extraction rms is 30.39, with extraction R of 0.46. The published external prediction rms of 16.00 is reached or improved upon in one of the 250 extractions. The published cross-validation rms of 19.70 is reached or improved upon in 14 of the 250 extractions. Both the cross-validation group and the external prediction group reported in 1998 were favorable draws.

On the larger set, training rms is 22.50 %HIA units, cross-validation rms is 25.56 and extraction rms is 25.77, with extraction R of 0.59. The same six descriptors therefore retain predictive information on the expanded set, while the gap between training and extraction error narrows.

### 3.2 Descriptor selection for the four deployed endpoints

The four searches each completed 250 extractions, absorption and the artificial-membrane peptides on a grid of 5 to 40 descriptors in steps of five, the Caco-2 small molecules on a grid of 5 to 55, and the Caco-2 peptides on a grid of 5 to 60. Averaged over the 250 sets each search selected, extraction rms and R are:

- absorption, rms 21.77 at R 0.73
- Caco-2 small molecules, rms 0.36 at R 0.84
- Caco-2 peptides, rms 0.53 at R 0.81
- artificial-membrane peptides, rms 0.83 at R 0.67

These values describe the searches; Table 3 reports the fixed consensus sets.

The number of descriptors settles inside the grid for three of the four endpoints (Figure 2). The Caco-2 small-molecule search selects the largest size its grid allows, where accuracy is flat in the descriptor count. The absorption search selects 25 descriptors most often, in 73 of 250 extractions, with 20 and 30 on either side; the artificial-membrane search also centers on 25, in 61 of 250. Extraction rms is nearly flat across the grid for both: for absorption it moves from 22.66 at 15 descriptors to 20.32 at 40, against a standard deviation near 1.80 within each count, and for the artificial-membrane peptides it stays between 0.82 and 0.85. Figure 2 plots the mean absolute error of the same extractions, which for absorption runs from 16.13 to 14.51 across the same range. The Caco-2 peptide search settles higher. On its grid of 5 to 60 it selects 45, 50 or 55 descriptors in 175 of 250 extractions, with a median of 50, and extraction rms stays between 0.52 and 0.55 from 35 to 60 descriptors. The search stops adding descriptors once the out-of-bag error no longer improves.

A small number of descriptors are chosen in most extractions, and the remainder are chosen far less often (Figure 3). For absorption, mean partial charge is chosen in all 250 extractions, hydrogen bond donor count in 72.40 percent, a carboxylic acid fragment in 62.00 percent, and topological polar surface area in 35.60 percent. For the artificial-membrane peptides, calculated logP is chosen most often, in 79.60 percent of extractions, and a term relating logP to polar surface area is chosen in 43.20 percent. For the Caco-2 peptides, Balaban’s J, a topological index, is chosen in 84.40 percent, followed by the minimum partial charge in 83.60 percent and the maximum in 79.60 percent.

**Figure 3.**
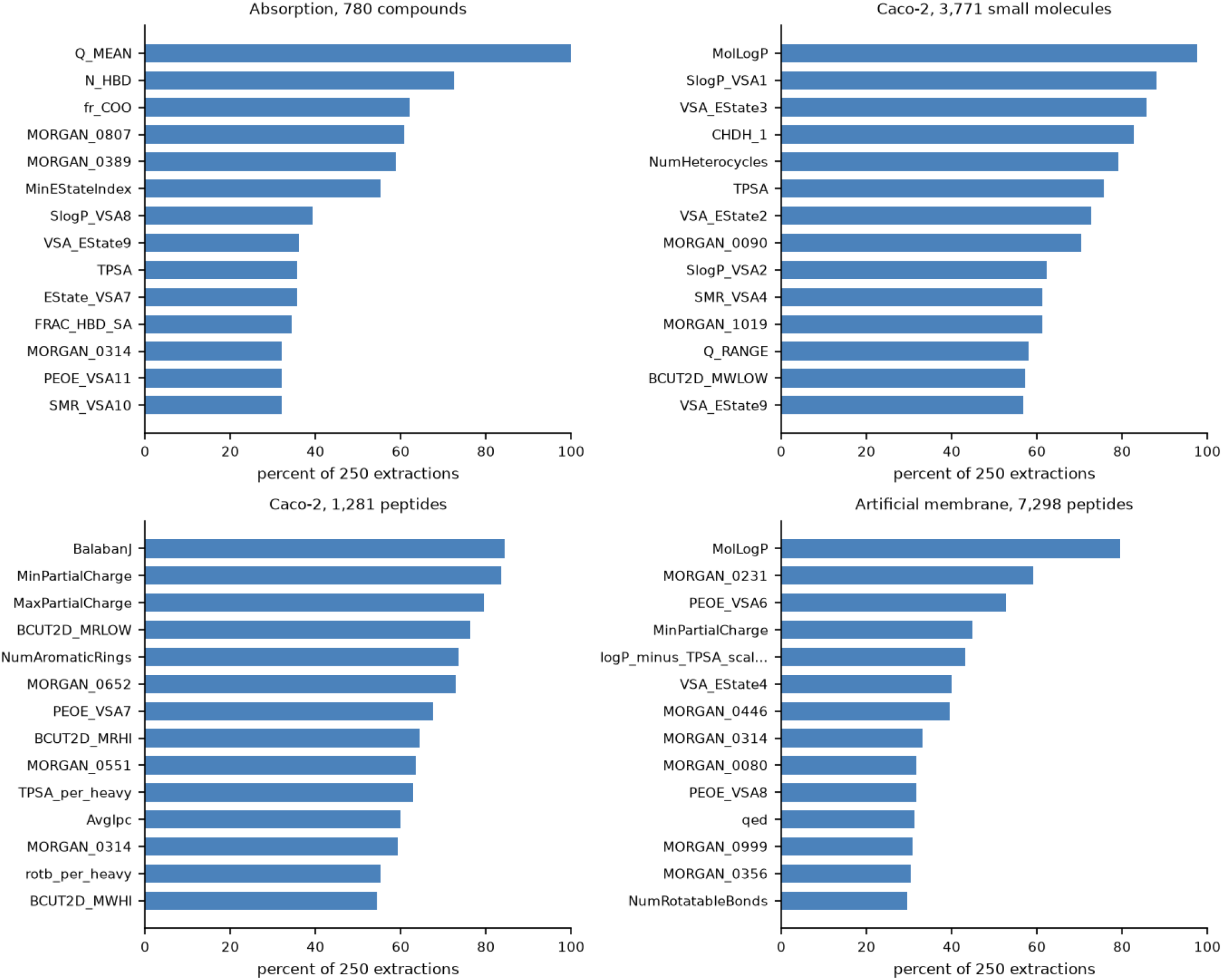
The fourteen descriptors most often chosen by the search, as a percentage of the 250 extractions that included them. Top left, absorption. Top right, the Caco-2 small molecules. Bottom left, the Caco-2 peptides. Bottom right, the artificial-membrane peptides. A descriptor appearing in every extraction is one the search never does without; the descriptors lower in each list are chosen less consistently. Charge, polar surface, charge-weighted surface and flexibility, the properties the 1998 descriptors measured, lead the Caco-2 peptide list as they do the absorption list, while calculated logP leads both the Caco-2 small-molecule and the artificial-membrane lists. The 1998 charge-on-donatable-hydrogens descriptor itself is the fourth most recurrent in the Caco-2 small-molecule search, chosen in 82.80 percent of extractions.

The last descriptor admitted to each consensus set is chosen about as often as the first one left out:

- absorption, 26.40 percent against 26.00
- Caco-2 small molecules, 32.40 against 31.20
- Caco-2 peptides, 37.20 against 36.80
- artificial-membrane peptides, 24.40 against 24.00

The descriptors chosen most often are therefore well determined, while those at the end of each set could be exchanged for others chosen nearly as often.

While the search was free to choose any of the six descriptors published in 1998, it chose only one of them often, a hydrogen bond acceptor surface term, in 29.20 percent of the absorption extractions, and that term is one of the 25 descriptors in the deployed absorption model. The properties those six descriptors measured were chosen repeatedly. Mean partial charge is chosen in every extraction, hydrogen bond donor count in 72.40 percent, and donor surface fraction in 34.40 percent. The search therefore replaces the individual 1998 descriptors and keeps the chemistry they measured.

Charge, polar surface, charge-weighted surface and flexibility are all still strongly selected, while the two three-dimensional properties, mass distribution and shape, are not.

The fixed consensus set is evaluated separately from the search that produced it. For absorption, it gives 20.47 rms at R 0.76 against 21.77 at R 0.73 for the search. For the Caco-2 small molecules the two agree, 0.35 against 0.36. For the Caco-2 peptides the consensus set gives 0.49 rms at R 0.84 against 0.53 at R 0.81 for the search. For the artificial-membrane peptides the two agree, 0.82 against 0.83.

The contribution of the descriptors can be separated from that of the model. Run under the same random forest as the deployed model, on the same compounds and the same extractions, the six descriptors published in 1998 give an extraction rms of 24.72 %HIA units at R 0.63, against 20.47 at R 0.76 for the 25 consensus descriptors. The 1998 network gives 25.77 at R 0.59 with those same six. Most of the improvement over the original model therefore comes from the descriptor pool and little from the model, and the six descriptors chosen in 1998 remain predictive on a set nine times the size of the one that produced them.

### 3.3 A small-molecule model for the Caco-2 assay

The deployed Caco-2 model is fitted on cyclic peptides, and it does not reach the accuracy of a model fitted on small molecules. Held fixed and scored on all 3,771 compounds, it reaches R 0.40 against measurement, with an rms of 0.63 log units against the 0.49 it achieves on peptides, and predicts a median 0.26 log units below measurement. Its own applicability domain places only 186 of the compounds, 4.93 percent, inside the region it was fitted in, and on those 186 it reaches R 0.27. While the peptide model carries real information onto these compounds, it sits outside its own applicability domain on 95.07 percent of them and reaches less than half the correlation it achieves on peptides. We therefore fitted a separate small-molecule model.

The same procedure was applied to the 3,771 compounds. The search selects 55 descriptors most often, in 128 of 250 extractions, with 50 in a further 98. The fixed consensus set gives an extraction rms of 0.35 log units at R 0.84 (Table 3).

Accuracy is flat in the number of descriptors on this endpoint. Evaluated on the extracted group of matched extractions, sets of 25 and of 100 descriptors give the same rms to within 0.01 log units, so the count the search settles on is not where the signal lies.

## 4. Prediction

Four models answer four distinct questions. The absorption model predicts the percentage of an oral dose absorbed through the human intestine. The Caco-2 small-molecule model predicts log10 Papp across a Caco-2 cell monolayer, the Caco-2 peptide model predicts the same quantity across the same monolayer for cyclic peptides, and the artificial-membrane model predicts log10 Papp across an artificial membrane. Permeability is expressed in cm/s before logarithmic transformation.

Each deployed model is fitted on every compound in its dataset using its fixed consensus descriptors. The absorption set contains 780 compounds, the Caco-2 small-molecule set 3,771, and the peptide sets 1,281 and 7,298. Fifteen percent of the compounds form the extracted group in each extraction, while the deployed model is fitted on all of them. Its reported rms, mae and R come from evaluating the fixed consensus set over the 250 extractions.

Absorption accuracy depends on where the compound sits. Pooled over the 29,250 predictions on extracted compounds of the fixed consensus set, rms varies with the band:

- below 30.00 percent absorbed, rms 33.61 on 4,000 predictions
- from 30.00 to below 80.00, rms 23.95 on 5,949 predictions
- at 80.00 and above, rms 15.11 on 19,301 predictions

The median absolute error is 10.08 %HIA units. Of those predictions, 53.74 percent fall within 11.00 units and 75.39 percent within 20.00. The model places a compound on the correct side of the 30.00 percent threshold in 92.64 percent of those predictions. Both the deployed model and the 1998 model over-predict poorly absorbed compounds, and the deployed model over-predicts them by less (Figure 4).

**Figure 4.**
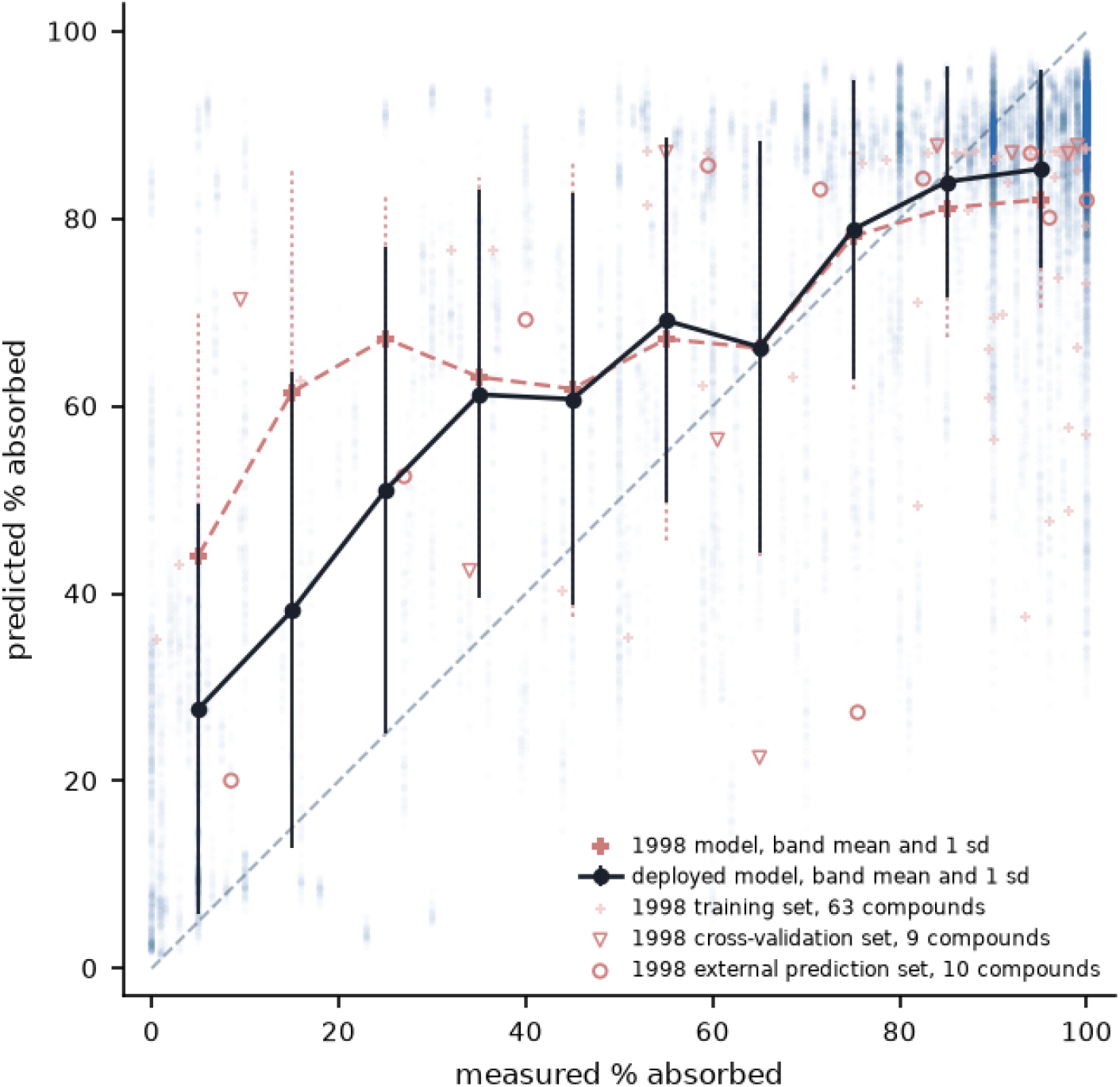
Predicted against measured absorption, for the deployed model and for the model it replaces. Each faint blue mark is one prediction on an extracted compound by the deployed model, which uses 25 consensus descriptors and a random forest; each of the 780 compounds falls in the extracted group about 38 times across the 250 extractions, giving 29,250 predictions. Black circles give the mean of those predictions within each band of ten % absorbed and the bars give one standard deviation. Red pluses with dotted bars give the same two quantities for the six descriptors published in 1998 under the 1998 network, run on the same compounds and the same extractions. Individual compounds of the 1998 set are drawn at their own mean prediction under that model, marked by the role they played in the original paper: thin pale pluses for the 63 of its 67 training compounds present here, open triangles for its 9 cross-validation compounds, and open circles for its 10 external prediction compounds. The dashed diagonal is perfect agreement. The two models agree above 55 % absorbed, and below 30 they separate, with the 1998 descriptors placing a compound near 44 % absorbed where it measures near zero and the consensus set placing it near 28

**Figure 5.**
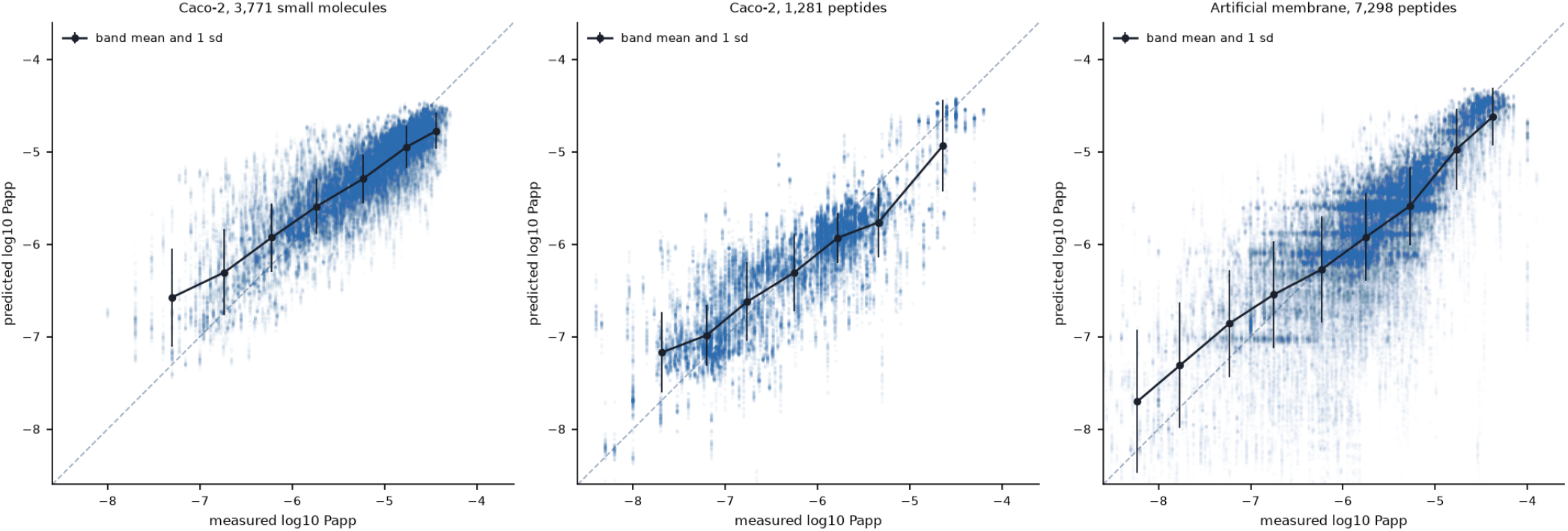
Predicted against measured permeability for the three deployed permeability models. Left, the Caco-2 small-molecule model, 55 consensus descriptors. Middle, the Caco-2 peptide model, 50 consensus descriptors. Right, the artificial-membrane model, 25 consensus descriptors. Each faint blue mark is one prediction on an extracted compound by the fixed consensus set over the 250 extractions. Black circles give the mean of those predictions within each band of half a log unit, placed at the mean measured value of the band, drawn where the band holds at least 50 compounds, with neighboring bands combined where together they reach 50, and the bars give one standard deviation. Peptides recorded at -10.00, below the detection limit, are not drawn; the small-molecule set contains no such values. The dashed diagonal is perfect agreement.

.For the Caco-2 small-molecule model, pooled over 142,000 predictions on extracted compounds, the median absolute error is 0.19 log units. Of those predictions, 68.21 percent fall within 0.30 log units and 91.55 percent within 0.60. The model places a compound on the correct side of the -6.00 log threshold in 90.87 percent of them. An rms of 0.35 log units corresponds to a factor of 2.25 in permeability, calculated from the unrounded rms.

For the Caco-2 peptide model, pooled over 48,000 predictions on extracted compounds, the median absolute error is 0.24 log units. Of those predictions, 59.55 percent fall within 0.30 log units and 85.58 percent within 0.60. The model places a peptide on the correct side of the -6.00 log threshold in 84.54 percent of them (Figure 5). An rms of 0.49 log units corresponds to a factor of 3.10 in permeability, calculated from the unrounded rms.

For the artificial-membrane model, pooled over 273,500 predictions on extracted compounds, the median absolute error is 0.29 log units. Of those predictions, 50.72 percent fall within 0.30 log units and 75.99 percent within 0.60. The model places a peptide on the correct side of the -6.00 log threshold in 79.40 percent of them. An rms of 0.82 log units corresponds to a factor of 6.68 in permeability, calculated from the unrounded rms.

All four models carry an applicability domain: the mean distance from a query to its five nearest training compounds in the model’s scaled descriptor space, with the 95th percentile of the same quantity over the training set as the threshold. That threshold is 0.55 for absorption, 0.84 for the Caco-2 small-molecule model, 1.10 for the Caco-2 peptide model and 0.33 for the artificial-membrane model.

The two peptide endpoints retain their assay identities. A Caco-2 measurement describes transport across a cell monolayer; an artificial-membrane measurement describes passive transport across a synthetic barrier. The endpoint used for a prediction is matched to the experimental question.

## 5. Discussion

The 1998 six-descriptor procedure can be reconstructed and applied repeatedly to its original compounds. Its training and cross-validation behavior must be distinguished from its external prediction. The published rms values were 9.40 for training, 19.70 for cross-validation and 16.00 for external prediction, in %HIA units.[1] With the published descriptors fixed across 250 stratified extractions, the corresponding means are 13.43, 29.37 and 30.39. Reproducing the procedure therefore provides a distribution of outcomes against which the original result can be judged.

The published external prediction result was a fortunate draw. Only one of the 250 extractions reaches 16.00 %HIA units or better. Repeated extraction turns the result of one draw into an estimate of expected performance on extracted compounds. The mean extraction R on the original set is 0.46.

The gap between training and extraction error was smaller on the larger absorption dataset. On the 86 compounds, the extraction rms of 30.39 is 2.26 times the training rms of 13.43. On the larger set, the same six descriptors give 22.50 for training and 25.77 for the extracted group, a ratio of 1.15. The descriptor set is unchanged. More compounds bring the error on training compounds and the error on extracted compounds closer together.

We separate descriptor search, consensus evaluation and deployment. Out-of-bag scoring keeps the search inside each extraction’s training group. The descriptors chosen most often then form one fixed consensus set, whose performance is measured directly over 250 extractions. The search covers 5 to 40 descriptors in steps of five, and extends to 60 for the Caco-2 peptides and 55 for the Caco-2 small molecules. The absorption and artificial-membrane searches both settle at a median of 25 descriptors, and their fixed consensus sets give an rms of 20.47 %HIA units and 0.82 log10 Papp units respectively. The Caco-2 small-molecule search settles at 55 descriptors with a fixed consensus set giving 0.35, and the Caco-2 peptide search at 50 with a fixed consensus set giving 0.49.

The practical result is four models fitted on every compound in their respective sets: percent human intestinal absorption, Caco-2 small-molecule permeability, Caco-2 peptide permeability and artificial-membrane peptide permeability. Each answers its own experimental question, and each is paired with the extraction statistics for its fixed consensus descriptors. These are the models provided through predicthia.ai.

## 6. Conclusion

The six descriptors published in 1998 give a mean extraction error over 250 extractions well above what the original single split reported, and the gap between fitting and prediction closes on the larger absorption set. Repeated extraction, an out-of-bag search inside each training group, and a fixed consensus set evaluated the same way are what separate a descriptor set’s behavior from the luck of one division of the compounds. On the larger absorption set the deployed model, 25 consensus descriptors fitted as a random forest, reaches an extraction rms of 20.47 %HIA units against 25.77 for the published six descriptors under the 1998 network. Four models built this way are in service at predicthia.ai, for percent human intestinal absorption, for small-molecule and peptide permeability across Caco-2 monolayers and for peptide permeability across an artificial membrane, each fitted on every compound in its dataset.

## 7. Data, Code and Availability

The absorption compilation[11] and CycPeptMPDB[10] are public. The completed reproduction results are recorded in holdout_A86fix.json, holdout_SIX781.json and holdout_SIX781RF.json, each containing 250 extractions, and Figure 1 is derived from those run artifacts. Structures for the larger absorption set were resolved from the PubChem identifiers provided by the compilation.

The four deployed models are named hia, caco2sm, pep_caco2 and pep_passive in the released files. Descriptor-count distributions and descriptor recurrence for all four endpoints are shown in Figures 2 and 3. The statistics of their fixed consensus sets are reported in Table 3 and recorded under frozen_consensus_set in hia_stats.json, caco2sm_stats.json, pep_caco2_stats.json and pep_passive_stats.json. All four endpoints are served at predicthia.ai, where a structure can be drawn or pasted and scored, and where the fitted models and the four datasets used to build them can be downloaded. The absorption set of 780 compounds, the 3,771 Caco-2 small molecules, the 1,281 Caco-2 peptides and the 7,298 artificial-membrane peptides are distributed as they were modeled, with their sources recorded alongside.

## Acknowledgments

We thank Peter C. Jurs and Jack W. Tolan, authors of the 1998 study this work revisits, for the original work on which it builds. We thank Emerson Hsieh for discussions related to the OpenADMET data, and the OpenADMET consortium and Expansion Therapeutics for releasing it openly.

